# Molecular maps of diseases from omics data and network embeddings

**DOI:** 10.1101/2025.11.25.689280

**Authors:** Dewei Hu, Anna-Lisa Schaap-Johansen, Julia Villarroel, Clara Ekebjærg, Simon Rasmussen, Daniel Hvidberg Hansen, Rasmus Wernersson, Lars Juhl Jensen

**Affiliations:** Novo Nordisk Foundation Center for Protein Research, Faculty of Health and Medical Sciences, University of Copenhagen, Copenhagen, Denmark; Novo Nordisk Foundation Center for Basic Metabolic Research, Faculty of Health and Medical Sciences, University of Copenhagen, Denmark; ZS Associates, ZS Discovery, Kgs. Lyngby, Denmark; Technical University of Denmark, Kgs. Lyngby, Denmark

## Abstract

Identifying disease-relevant proteins and pathways remains a fundamental challenge in understanding disease mechanisms and supporting therapeutic development. While omics analyses can provide valuable insights, they typically consider each gene/protein separately rather than at the level of biological systems. This can be addressed by combining the omics data with protein networks. We integrate disease-specific omics data with a universal functional association network from STRING, which we represent using node2vec embedding. This way, we constructed disease maps for seven diseases spanning inflammatory, oncological, neurological, and vascular diseases based on genetics, transcriptomics, somatic mutation, and proteomics data. Compared to omics analysis alone, the use of a simple linear model on top of network embedding enabled us to identify 2–4 times as many known disease-relevant proteins at the same specificity. Clustering of the resulting disease maps revealed both functional modules shared by many diseases, such as inflammatory pathways and cancer hallmarks, and disease-specific modules, such as keratinization in atopic dermatitis and extracellular matrix remodeling in aortic aneurysm. Together, these results highlight the value of protein network embedding when analyzing omics data to understand diseases.

## Introduction

Mapping the molecular basis for human diseases is important for understanding them better and for identifying biomarkers and potential new drug targets. The many new omics technologies have enabled systematic identification of disease-associated genes and proteins, including genome-wide association studies (GWAS) ^1^, transcriptomics^2^, somatic mutations^3^, and proteomics^4^. These modalities provide complementary views of a biological system, each identifying a subset of protein-coding genes involved in a disease. However, these omics data types do not directly capture the interplay of the proteins in complexes, pathways, and regulatory networks.

Protein–protein interaction (PPI) networks capture physical and functional relationships between proteins that are essential for understanding biological processes^5^. For example, the STRING database^6^ is one of the most comprehensive resources for functional associations between proteins, including evidence from experimental data, text mining, and curated knowledge from other databases into a graph structure. Systematic integration of omics data and PPI networks can map disease-associated proteins within a broader functional context^7^, effectively going from targeting individual molecular hits to functionally connected protein modules.

Network embedding is a method that converts nodes in a graph into low-dimensional vector representations, which are useful for downstream network analyses^8^. In bioinformatics, network embedding has been used to capture the functional relationships in PPI networks^9^. Many algorithms have been proposed for network embedding, including deepwalk^10^, node2vec^11^, and graph neural networks^12^. For PPI networks, weighted node2vec has been demonstrated to be especially well-suited, capturing functional associations more accurately than alternative graph neural network approaches^13^.

In this study, we combine omics data with network embeddings to create molecular maps of diseases. We labeled proteins as associated or not associated with a given disease based on disease-specific omics data and trained a logistic regression model to predict disease association from a node2vec embedding of STRING. We performed analysis across seven diseases, including inflammatory, oncological, neurological, and vascular diseases. We demonstrate that this network-based AI method identifies 2–4 times as many known disease-relevant proteins as the omics data alone at the same specificity. We further demonstrate that this integration analysis captures meaningful functional modules by clustering the disease-associated proteins.

## Results

### Enhanced disease protein mapping through a network-based AI method

To move beyond the limitations of traditional omics-based analysis for mapping disease-associated proteins, we explored whether applying a network-based AI method, by integrating omics data with protein-protein interaction networks, could enhance the disease-relevant protein identification. We generated 64-dimensional node2vec embeddings from the STRING network to represent the 19,622 proteins as feature vectors, which we use as the foundation for developing disease-specific models. Disease-specific omics data (e.g., GWAS and transcriptomics) were used to assign positive and negative labels to proteins for each disease of interest. We trained logistic regression models using the embeddings and evaluated performance against literature-derived gold standards. The top predicted disease-associated proteins were inspected using network clustering and enrichment analysis.

Seven diseases were selected for evaluation, including atopic dermatitis, ulcerative colitis, focal epilepsy, colorectal adenocarcinoma (referred to as colorectal cancer), diffuse large B-cell lymphoma (referred to as lymphoma), melanoma, and aortic aneurysm. These diseases were selected based on the following criteria: high-quality omics data are available, the diseases are prevalent and of significant interest to industry practitioners, each disease affects a clearly defined organ or tissue, and omics data can be obtained from ante-mortem tissue. The last criterion is specifically why we selected focal epilepsy over other diseases of the central nervous system, as the RNA-seq data can be collected from surgically resected brain tissue.

We benchmarked our approach using five-fold cross-validation and an independent literature-based gold standard. The gold standard was constructed from the DISEASES database^14^, which collects text-mined protein-disease associations from the biomedical literature. To ensure fair evaluation, we balanced the studiedness of the proteins labelled as positive and negative examples (see Methods). When assessing the omics-based disease-protein association against the gold standard, we ranked proteins by their p-values from the statistical analyses when available. In datasets with no p-values, all disease-associated proteins were considered as an unranked list, and thus a single point was evaluated rather than a full-ranking curve.

For inflammatory diseases (Fig 2A and Fig 2B), the network-based AI method was significantly better than omics alone analysis (DeLong test^15,16^ P < 10^-8^ in atopic dermatitis and P < 10^-16^ in ulcerative colitis, see Table S4 and Table S5), and achieved 2–4 times higher true positive rates at a fixed 5% false positive rate compared to omics-only analysis. Similar gains were observed in focal epilepsy (Fig 2C), aortic aneurysm (Fig 2D), and cancers (Fig 3A). These results suggested that integrating protein network context can enhance disease-associated protein mapping. Cross-validation analysis showed consistent performance across the five folds for each disease (Fig 3B and Fig S1), showing the robustness of the network-enhanced approach.

**Fig 1.**
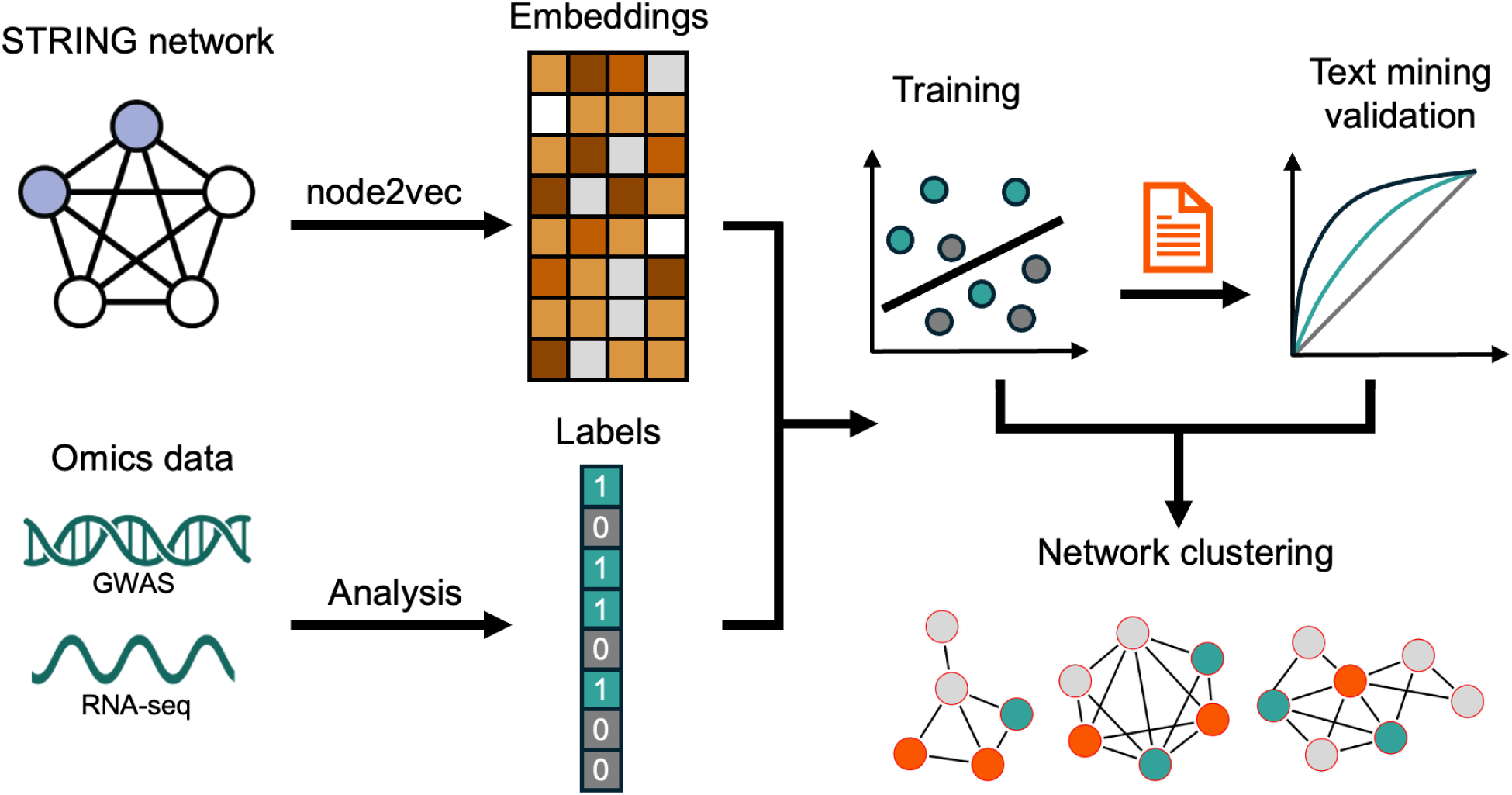
Analysis workflow of the network-based method for disease protein mapping. The proteins in the STRING protein-protein interaction network are represented as embeddings using the node2vec algorithm. Omics datasets are analyzed to generate binary labels indicating disease association for each protein. These protein embeddings and labels are used to train logistic regression models for disease prediction, with model performance validated against text-mining-derived associations. Finally, network clustering is applied to highly predicted disease proteins to identify functional modules and pathways underlying disease mechanisms.

**Fig 2.**
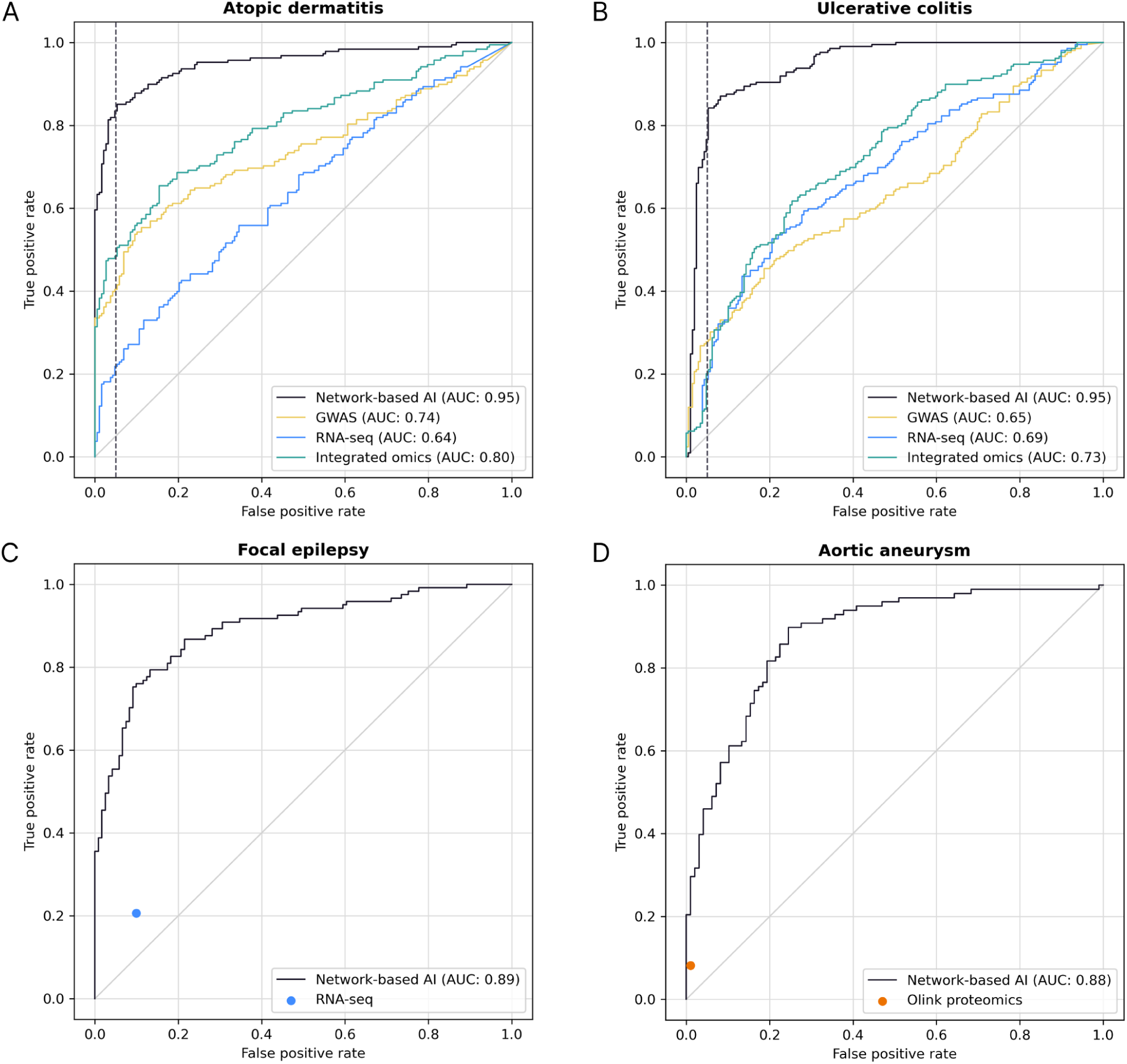
Literature-based benchmark results for non-cancer diseases. **(A)** Atopic dermatitis. **(B)** Ulcerative colitis. **(C)** Focal epilepsy. **(D)** Aortic aneurysm. In **(A)** and **(B)**, we present the ROC curves and AUC scores for both the omics-only analysis and the network-based AI method. In **(C)** and **(D)**, we use a dot to indicate the true positive rates and false positive rates achieved by the omics-based analysis. The performance of the network-based AI method is shown in black in all panels.

**Fig 3.**
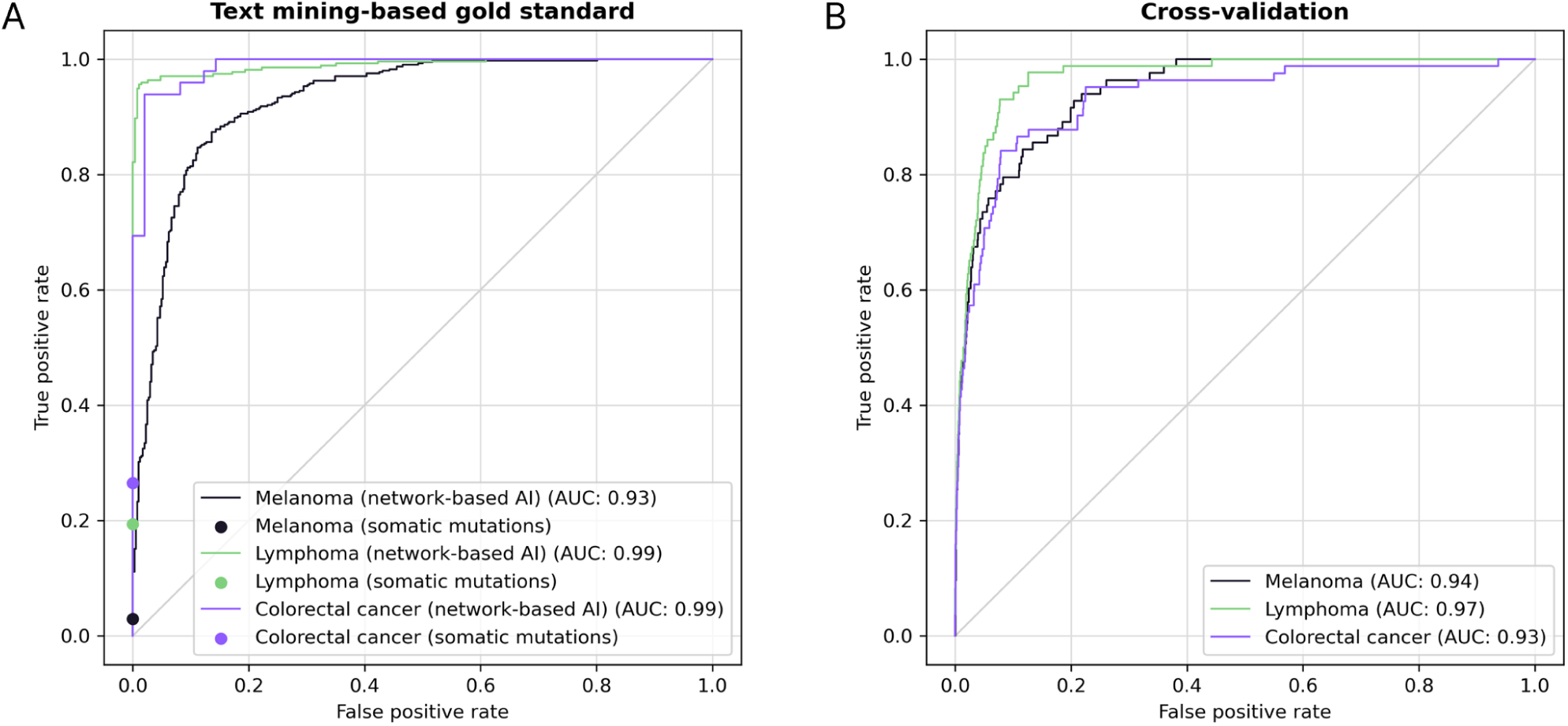
ROC curves for three cancers. **(A)** Benchmark against the literature-derived gold standards. We use curves to illustrate the performance of the network-based AI method, and dots to indicate the true positive rates and false positive rates achieved by the omics-based analysis. **(B)** Aggregated cross-validation ROC curves. The ROC curves show the robustness of the network-based AI method.

Together, these findings demonstrate that incorporating PPI context through network embeddings substantially enhances the identification of disease-relevant proteins compared with omics data alone. The consistent performance across diverse diseases and validation folds underscores that this network-based AI method can provide a robust and generalizable disease protein mapping.

### Shared inflammatory pathways and disease-specific signatures

To investigate how the network-based AI method captured inflammatory disease mechanisms, we analyzed predictions for atopic dermatitis and ulcerative colitis. We selected the top predicted proteins (1,717 in atopic dermatitis and 2,647 in ulcerative colitis) using a 5% false positive rate threshold, defined against the literature-based gold standard. For these two sets of proteins, we retrieved the high-confidence STRING functional association networks and identified functional modules through MCL clustering^17^ (see Methods). Shared inflammatory mechanisms were evident between the two diseases, as reflected by the substantial overlap of disease-associated proteins (Fig 4D) and with the inflammatome dataset.

**Fig 4.**
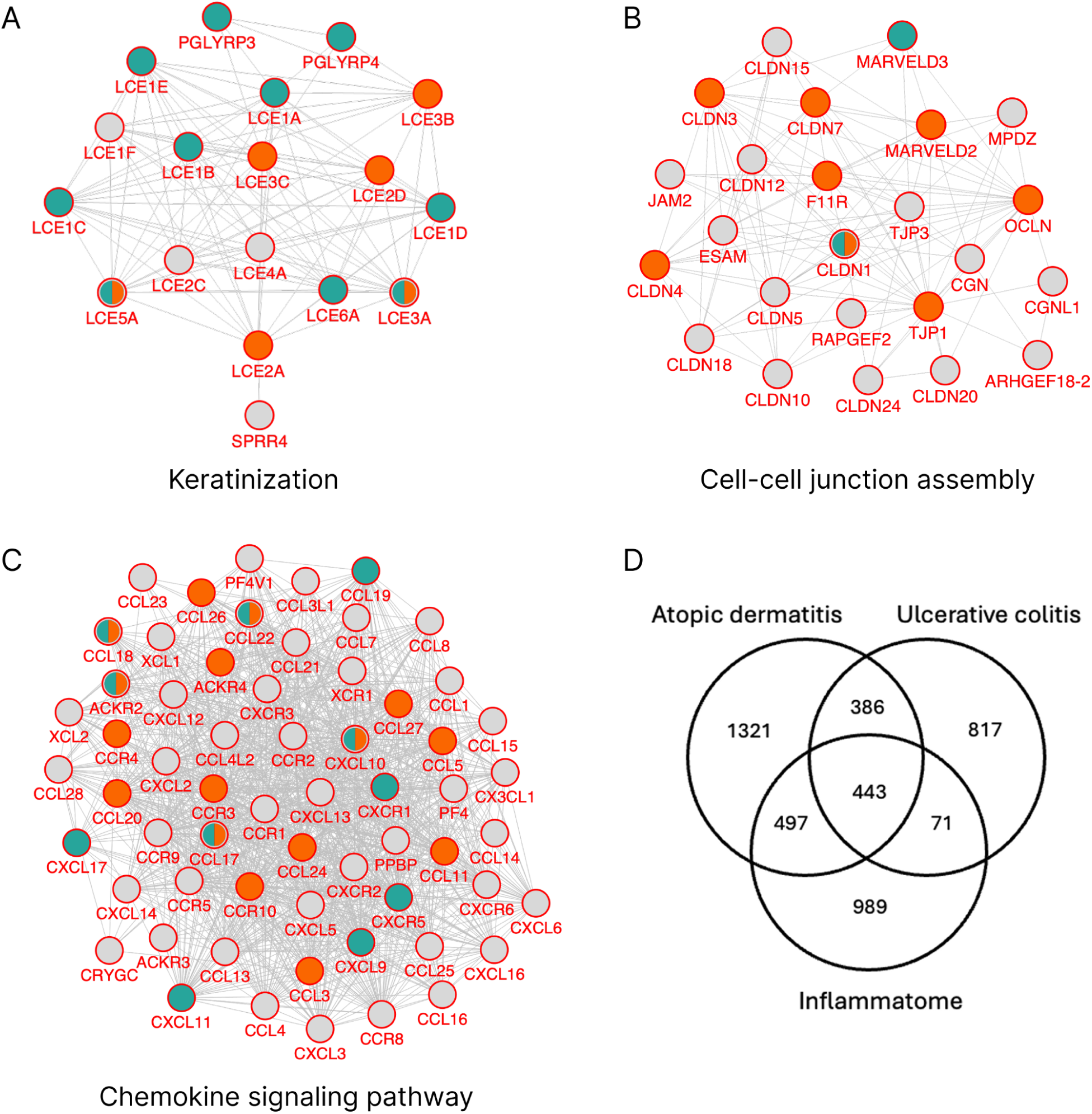
Network modules and Venn diagram of inflammatory diseases. The network modules contain gold-standard proteins (orange), proteins highly ranked from integrated omics analysis (teal), and novel candidates (light gray). (A) Proteins involved in keratinization in atopic dermatitis. (B) Proteins involved in cell-cell junction assembly in ulcerative colitis. (C) Chemokine signaling is prominent in both atopic dermatitis (shown here) and ulcerative colitis. (D) Venn diagram showing the overlaps between the proteins predicted to be involved in atopic dermatitis and ulcerative colitis, and proteins that are part of the inflammatome^21^.

To characterize the functional modules underlying each disease, we performed functional enrichment analysis on the identified network clusters. The atopic dermatitis network contained a disease-specific module related to keratinization (Fig 4A), reflecting the skin barrier dysfunction characteristic of the disease. Keratinocytes form the physical skin barrier and facilitate communication between innate and adaptive immune responses, and the damaged skin barrier can activate the keratinocytes, among the dysregulation of immune responses^18^.

The ulcerative colitis network similarly contained a disease-specific module, this time related to cell-cell junction assembly (Fig 4B), a process by which specialized protein structures form connections between neighboring cells. This helps maintain structural integrity, enables communication, and regulates transport across tissues. In the intestine, the bicellular tight junction plays an essential role in forming the physical barrier, and its dysfunction can lead to diseases such as ulcerative colitis ^19^. The claudins in this functional module are used as biomarkers for ulcerative colitis^20^.

In both diseases, the largest clusters were related to general inflammation, exemplified by the chemokine signalling module in the atopic dermatitis network (Fig 4C). Comparison with the inflammatome ^21^, a consensus set of proteins commonly regulated across inflammatory diseases, revealed that 36% of the proteins in atopic dermatitis and 30% in ulcerative colitis overlapped with this shared inflammatory signature (Fig 4D). Thus, one-third of the altered proteins in each disease reflected a general inflammatory response observed across multiple conditions, while the remaining 60-70% were disease-specific, indicating that both diseases shared a core inflammatory program alongside their distinct molecular characteristics.

Our analysis identifies a cohesive inflammatory core that underlies two chronic diseases, atopic dermatitis and ulcerative colitis. Each condition retains a distinct molecular signature reflecting its tissue-specific context. By comparing these patterns, we demonstrate that systemic inflammation arises from both common and condition-specific mechanisms, pointing to potential cross-disease intervention points.

### Neurological mechanisms and immune response in focal epilepsy

In neurology, a major challenge in disease protein mapping is the reliance on post-mortem brain samples. These tissues often contain confounding signals from degradation and late-stage pathology, obscuring the molecular changes that drive disease onset^22^. To demonstrate the applicability of our approach within neurology, we thus focused on focal epilepsy, a condition with available transcriptomics data from living patients^23^. These datasets provide a rare window into disease-related alterations without the confounding factors inherent to postmortem material. Compared to the transcriptomics-based statistical analysis, the network-based AI approach expanded the disease mapping to cover nearly 4 times as many proteins associated with focal epilepsy in the literature (Fig 2C).

Functional enrichment analysis of the predicted disease-associated proteins validated their relevance to the disease, identifying key pathogenic mechanisms that align with established focal epilepsy pathophysiology. This includes several Gene Ontology (GO) terms related to synaptic mechanisms, such as synaptic transmission, postsynapse organization, glutamate signalling pathway, and potassium and sodium transportation (FDR < 10^-5^). We also observed enrichment for terms related to immune response, such as T cell activation and peptide antigen assembly with MHC class II complex (FDR < 10^-4^).

In summary, these results indicate that the network-based AI method can capture both neuronal signaling and immune processes central to focal epilepsy, expanding beyond the scope of transcriptomic analysis.

### Cancer hallmarks and tumor type-specific functional modules

Cancers were analyzed because they remain among the most prevalent and deadly diseases worldwide, posing major challenges for diagnosis and treatment. Understanding their molecular mechanisms is key to advancing precision medicine. From a systems perspective, cancers are also ideal for testing the network-based AI method, as they combine shared hallmark processes with distinct tumor-type–specific regulatory programs.

We evaluated the method on colorectal cancer, lymphoma, and melanoma. These cancers each have approximately 80 associated proteins in the training data, with minimal overlap between each other (Jaccard index < 0.14, Table S3). Despite this low initial overlap in genes with somatic mutations, the network-based AI predictions identified 1,388 proteins shared across all three cancers (Fig 5A).

**Fig 5.**
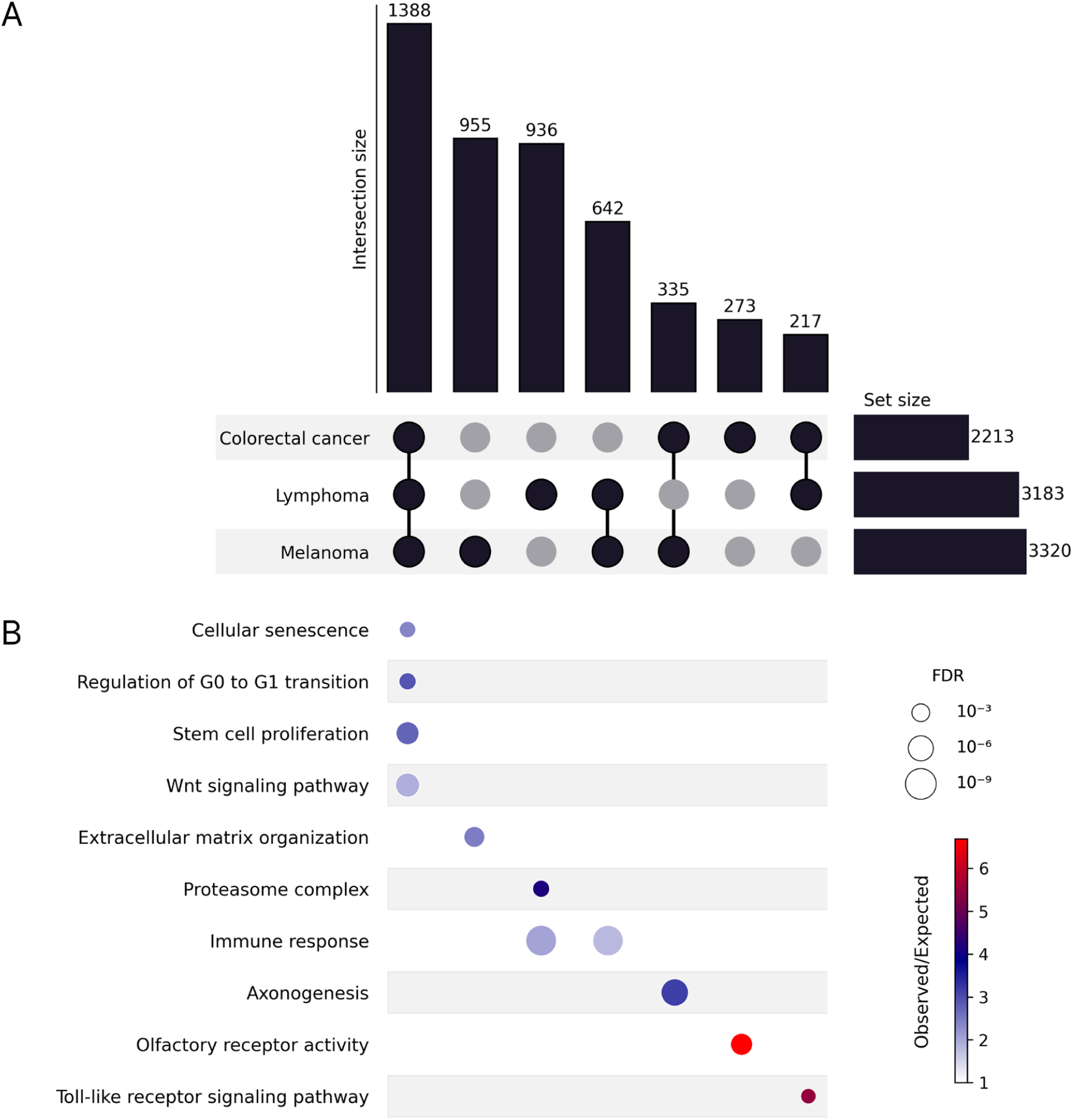
Analysis of disease-associated proteins in three cancers. **(A)** UpSet plot showing the intersection of gene sets associated with colorectal cancer, lymphoma, and melanoma. Horizontal bars represent the total number of genes identified in each cancer, and vertical bars indicate the size of intersections among the sets as defined by the connected sets or single sets. **(B)** Bubble plot of functional enrichment analysis of each gene set intersection (corresponding to the columns in panel **(A)**. Bubble size represents the false discovery rate of the processes, and color intensity corresponds to the ratio of the number of proteins observed against the expected in a given Gene Ontology term.

To characterize what was common and unique to the cancers, we analyzed which GO terms were enriched for the proteins that were shared among the cancers or unique to each (Fig 5B). As expected, the proteins common to all three cancers were predominantly involved in core cancer processes, such as cellular senescence, cell cycle regulation of G0 to G1 transition, stem cell proliferation, and Wnt signaling. Immune response and toll-like receptor signaling were enriched in the lymphoma–melanoma and colorectal cancer–lymphoma intersections, respectively. The proteins common to colorectal cancer and melanoma showed enrichment for axonogenesis, consistent with reports linking neural infiltration to cancer prognosis in solid tumors, including colon and skin cancer^24^.

Each cancer type further exhibited distinct functional signatures. Lymphoma-specific proteins were enriched for immune response, reflecting the B-cell origin, and for the proteasome, consistent with the use of proteasome inhibitors in lymphoma treatment^25,26^. Melanoma-specific proteins showed enrichment for extracellular matrix organization, which agrees with the role of matrix degradation in melanoma migration, invasion, and metastasis^27^.

In colorectal cancer, we observed a surprising enrichment for olfactory receptors. Since olfactory receptors constitute one of the largest gene families in the human genome, the enrichment analysis (Fig 5B) was performed using the entire human proteome as background, ensuring that this did not occur by chance. Moreover, several studies have shown that olfactory receptors play a role in colorectal cancer, including OR51B4, OR7C1, TAARs, FPRs, and MS4As^28–30^. Although these specific olfactory receptors were not identified in our analysis, they support the idea that other olfactory receptors may also be involved in colorectal cancer, in which case they could be potential drug targets^30^.

The network-based AI method revealed a large shared core of cancer-associated proteins, representing universal hallmarks such as cell cycle control, senescence, and Wnt signaling, while preserving distinct functional modules that reflect each tumor’s biology. This demonstrates that integrating network context can uncover both the conserved molecular foundations of cancer and tumor-type–specific pathways with therapeutic potential.

### Extending predictions beyond platform coverage limitations in aortic aneurysm

Aortic aneurysm was selected as a case study to test whether the network-based AI method can extend disease protein mapping beyond the limited coverage of targeted omics platforms. This condition exemplifies a challenge in translational research that key disease mechanisms might involve proteins not detectable by standard assays.

To illustrate this, we applied the network-based AI method to Olink proteomics data (2,187 measured proteins, 109 positives) from aortic aneurysm samples, resulting in 996 predicted disease-associated proteins. Of these proteins, 271 were measured but not identified as differentially regulated, whereas 658 were not measured by the Olink platform at all. To better understand the biological context of these predicted proteins, we performed a network analysis, which revealed two key functional modules related to the extracellular matrix (Fig 6). The extracellular matrix provides structural support^31^ and elastic recovery for soft tissues undergoing continuous mechanical stress^32^. Weakening of it can thus reduce aortic wall elasticity^33^ and thereby lead to aortic aneurysm^34^.

**Fig 6.**
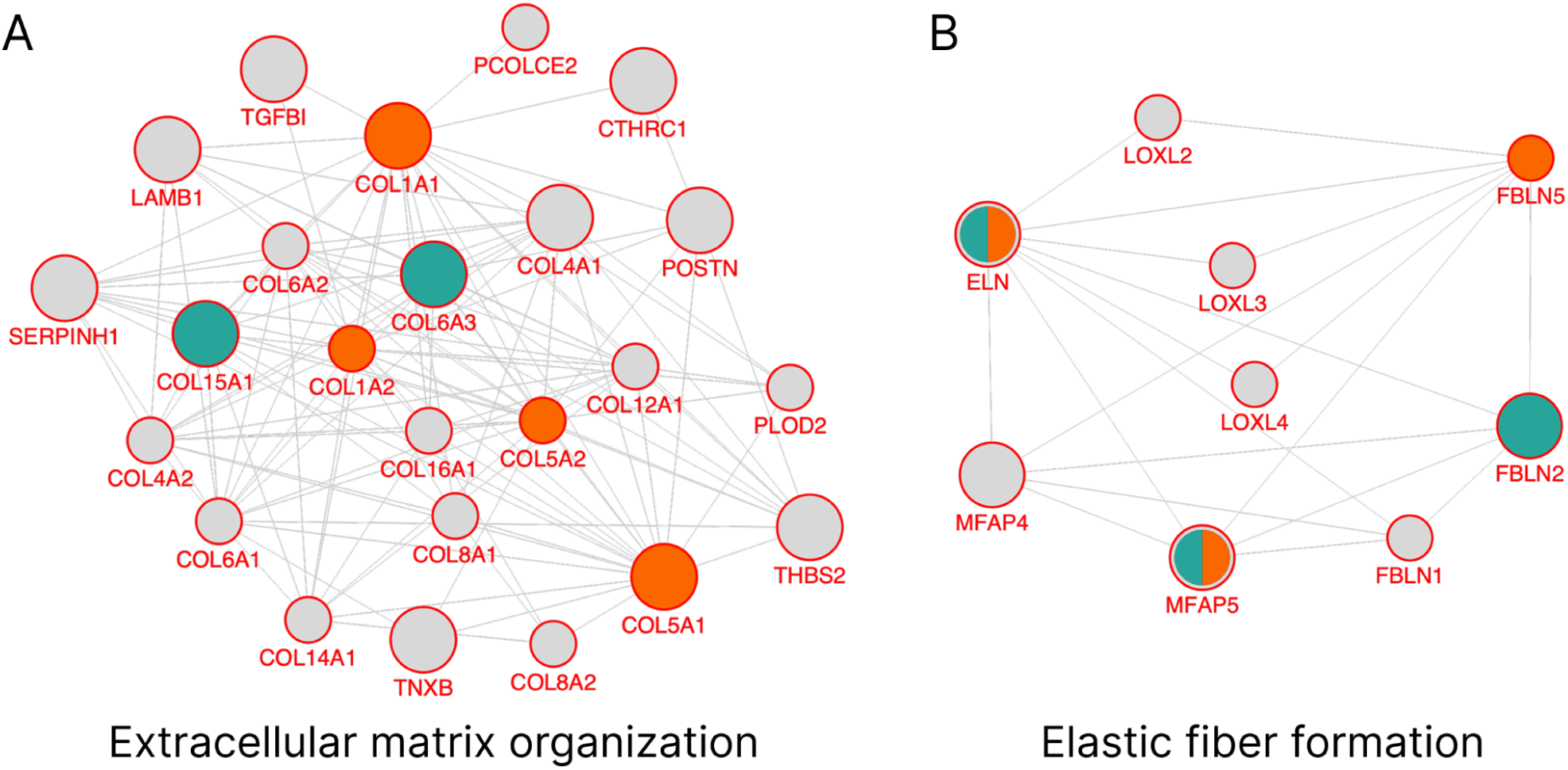
Network modules of aortic aneurysm-associated proteins. The network modules contain gold-standard proteins (orange), proteins highly ranked from integrated omics analysis (teal), and novel candidates (light gray). Large nodes represent proteins measured in proteomics data, while small nodes indicate unmeasured proteins. **(A)** Extracellular matrix organization provides physical support for organs and tissues. **(B)** Elastic fiber formation. Elastic fibers, as components of the extracellular matrix, provide tissues with elasticity and resilience.

These results show that the network-based AI method can expand disease protein predictions beyond experimentally measured targets, effectively bridging gaps caused by limited assay coverage. By integrating network information, the approach highlights biologically plausible candidates that are inaccessible to direct proteomic profiling, illustrating its value for uncovering hidden disease mechanisms.

## Discussion

In this work, we show how embedding of PPI networks can be used to construct molecular maps for a diverse set of diseases, using multiple omics modalities. We demonstrate that compared to omics data alone, network-based AI can identify more than twice as many of the disease-related proteins found in literature at the same false positive rate. This also enables looking beyond the proteins measured by targeted omics platforms, such as Olink proteomics. Just as important, clustering of the resulting protein networks groups these proteins into functional modules that capture the unique features of each disease as well as shared patterns both in inflammatory diseases and in oncology.

It is important to acknowledge that no benchmark is without flaws, and any gold standard derived from literature will unavoidably be subject to study bias. While we have done our best to mitigate this by balancing our gold standards to make sure that positive and negative examples are approximately equally well studied, this still results in a gold standard predominantly composed of well-studied proteins. As understudied proteins will have many fewer interactions in STRING than the gold-standard proteins, one should expect any network-based approach to perform worse on understudied proteins.

The problem of study bias can be avoided by relying on systematic omics data rather than literature. Since we use omics-based labels for training to avoid building study bias into our models, benchmarking on omics was done in the form of cross-validation. The results are more modest than what we see for the literature-based gold standard, but there is good reason to believe that cross-validation underestimates the true performance. The premise of our work is that each omics modality will identify only a subset of the proteins relevant to a disease, and that PPIs can bring in the missing proteins. Counting all the latter as false positives is thus bound to put any network-based approach in a bad light.

Despite these caveats, our analysis demonstrates the utility of network embeddings in mapping disease–protein associations. The molecular maps can both help improve disease understanding and serve as a first step in therapeutic target discovery, beginning with a broad spectrum of candidate proteins and progressively narrowing toward those with true potential for pharmacological intervention. We believe that this analysis provides valuable insights for future target identification and validation efforts.

## Methods

We applied a network-based AI method to predict disease-associated proteins by integrating omics-derived signals with protein-protein functional interaction networks. Specifically, we generated protein embeddings using node2vec on the STRING functional network. For each disease, omics data were used to define positive and negative sets, which served as training samples for a logistic regression model. The trained model then predicted the probability of each human gene being associated with the disease. Model performance was evaluated using five-fold cross-validation and validated against an independent literature-based gold standard.

We benchmarked the network-based approach across seven diseases representing diverse pathological contexts and omics modalities. Our analysis included inflammatory diseases (atopic dermatitis and ulcerative colitis using integrated GWAS and transcriptomics data), cancers (colorectal adenocarcinoma, diffuse large B-cell lymphoma, and melanoma using somatic mutation data), neurological disorders (focal epilepsy using transcriptomics), and vascular disease (aortic aneurysm using proteomics data). We describe how we processed each type of omics data below.

### GWAS data

GWAS summary statistics for atopic dermatitis and ulcerative colitis were obtained from publicly available resources. For atopic dermatitis, the dataset GCST90244788 was downloaded from the EMBL-EBI GWAS catalog^35^. Ulcerative colitis summary statistics were obtained from two independent sources: the FinnGen (release 11)^36^ and a previously published GWAS study from 2015^37^. From the 2015 GWAS study, we used EUR.UC.gwas_info03_filtered.assoc with summary statistics in Europeans. The FinnGen file was filtered by excluding variants with a minor allele frequency <1%. Gene-level analysis was performed using MAGMA^38^ for all three datasets. The results from MAGMA were mapped from Entrez gene IDs to STRING IDs via Ensembl and UniProt. For ulcerative colitis, results from the FinnGen and 2015 GWAs datasets were further combined using Fisher’s method to obtain a joint gene-level significance score.

### Transcriptomics data

Transcriptomics data was downloaded from Gene Expression Omnibus^39^, for atopic dermatitis (GSE121212) and ulcerative colitis (GSE66407, GSE109142, and GSE166925). DESeq2^40^ was used to test for differential expression between disease and control.

### Somatic mutation data

For each of the three cancer types, we downloaded the mutational cancer driver genes from IntOGen^41^. The number of associated proteins is shown in Table S1. We used all proteins associated with a cancer as positive samples for that cancer, and all other proteins as negatives. The number of positive and negative samples for each cancer is shown in Table S1.

### Olink proteomics data

The proteomics data and analysis were sourced from Olink Insight of all UK Biobank phenotypes (https://insight.olink.com/olink-data/ukb-diseases), where each protein was associated with phenotypes (including diseases) by hazard ratios and p-values. In total, 2,922 proteins were measured using Olink assays, organized by panels (cardiometabolic, inflammation, neurology, and oncology). We define positives as proteins with a hazard ratio > 2 and p-value < 0.01 for aortic aneurysm, and negatives as proteins that were measured but had neither a hazard ratio > 2 nor a p-value < 0.01. The oncology panel contained very few positives compared to the other panels, and we thus based all analyses on the 2,190 proteins from the three other panels. The number of positive and negative samples is shown in Table S1.

### Integration of omics modalities

For both inflammatory diseases, we integrated the GWAS and transcriptomics data. We obtained the omics datasets mentioned before from public sources for each disease and analyzed them using standard statistical methods to obtain a p-value for each protein. The omics datasets for each disease were integrated by using Fisher’s method^42^ to combine the GWAS and transcriptomics p-values into a single p-value for each protein. We obtained positive examples for training the model from the integrated omics data by ranking proteins based on the integrated p-values and selecting the proteins with the highest ranking. For atopic dermatitis, we selected the top 1,000 proteins, and for ulcerative colitis, we selected the top 1,300 proteins based on ROC curves against the literature-based gold standard (Fig 2A and Fig 2B). Negative samples were drawn from the complement of the expanded positive set, which consisted of all proteins excluding those ranked within the top 4*N positions, where N represented the number of top-ranked proteins initially selected. The number of positive and negative samples for each inflammatory disease is shown in Table S1.

### Network embedding of the STRING network

We downloaded the functional association network of all human proteins from the STRING database^6^ (version 12.0). We created a 64-dimensional embedding of this network using the PecanPy^43^ implementation of the node2vec^11^ algorithm, taking into account the confidence score of each edge. Except for the dimensionality of the embedding, we optimized the hyperparameters of node2vec as described elsewhere^44^. That work systematically tuned the walk length, number of walks per node, and the return (p) and in–out (q) parameters to maximize the functional coherence of neighboring proteins. In this study, we tried to make embedding dimensions as low as possible. As in many diseases, there are not too many positive samples. In this case, we searched for the best hyperparameters in a setting where the embeddings could be 32, 64, or 128, but extended the exploration by testing more walks per node (50, 70, 100) and longer training epochs to ensure convergence on the larger STRING network. The best-performing configuration (p = 0.1, q = 0.9, 100 walk length, 100 walks per node, 10 epochs, and 64 dimensions) was selected based on the link prediction standard mentioned by the same research^44^.

### Logistic regression model training

For each disease and data type, we first performed a five-fold cross-validation on the training set using a logistic regression model. We then trained a logistic regression model on the full training set and validated it using the literature mining test set. All the training and tests were implemented with scikit-learn^45^.

### Gold standard from literature mining

We evaluated the predictive performance of each trained model using a test set (Table S2) based on protein–disease associations obtained from text mining of the scientific literature (DISEASES^14^). Positive examples were required to have a text-mining confidence score of at least 2.0 for the disease in question, and negative examples were not allowed to be associated. The test set was corrected for study bias to ensure that model performance was not driven by differential publication frequency. For each positive example (disease-associated protein), we selected a negative example (disease-unrelated protein) with a similar level of literature coverage (a number of PubMed mentions within a factor of two of its paired positive). This yielded a balanced dataset in which positive and negative proteins were equally studied, preventing the model from exploiting literature bias.

### Prediction of disease-associated proteins

Using the trained logistic regression models within *the network-based AI method*, we predicted association probabilities for the entire proteome. For each disease, we applied a probability threshold corresponding to a 5% false positive rate and identified proteins below this threshold as mapped disease-associated proteins. A 5% false positive rate was chosen as the operating point to balance sensitivity and specificity, and to ensure robust recovery of known disease-associated proteins.

### Network analysis of top disease predictions and function enrichment analysis

For inflammatory diseases and aortic aneurysm, we created network visualizations by retrieving functional association networks from STRING (confidence ≥ 0.7) using Cytoscape^46^ (version 3.10.3) stringApp ^47^ (version 2.2.0). Each disease network was clustered using the MCL clustering algorithm^17^ (inflation value = 4). The network modules were functionally annotated using enrichment analysis, testing which biological terms were significantly overrepresented in each module using the whole human proteome as the statistical background. False discovery rate was assessed using a hypergeometric test with each category (Reactome, WikiPathways pathways, as well as Gene Ontology biological processes, molecular function, and cellular component) using the Benjamini–Hochberg procedure, and terms with FDR < 0.05 were considered significant. When analysing protein functions that were shared or unique among different cancers (Fig 5B), we used the union set of proteins in three cancers as background, and performed the enrichment analysis for each subset.

## Availability

The source code, training and evaluation data, and supplementary files are available at https://github.com/larsjuhljensen/net2rank and https://zenodo.org/records/16919169. The license does not permit us to redistribute the Olink data for aortic aneurysm, but it can be downloaded at https://insight.olink.com/olink-data/ukb-diseases.

## Author contributions

L.J.J. conceptualized the study and developed the scientific framework. D.H., A-L.S-J., J.V., and C.E. jointly collected the omics data. D.H., A-L.S-J., J.V., and C.E. processed the data. D.H. created the network embeddings. L.J.J. created the text mining benchmark dataset. D.H. and A-L.S-J. performed machine learning. J.V. performed the network analysis. D.H.H. generated all ROC curves and associated statistical evaluations. D.H., A-L.S-J., and L.J.J. drafted the initial manuscript. S.R. provided feedback on the analysis and manuscript. All authors reviewed the manuscript, provided revisions and feedback, and approved the final version.

## Conflict of interest

S.R. is the founder and owner of the Danish company BioAI and has performed consulting for Sidera Bio ApS. L.J.J., D.H.H., A-L.S-J., and J.V. are employees of ZS Associates. The study was conducted as part of ZS Discovery’s internal research activities. The authors declare no other competing financial interests.

## Funding

D.H. was supported by the Novo Nordisk Foundation (grants NNF14CC0001 and NNF20SA0035590). S.R. was supported by the Novo Nordisk Foundation (grant NNF23SA0084103).

## Notes

https://github.com/larsjuhljensen/net2rank

https://doi.org/10.5281/zenodo.16919169

