## Supplementary file for "Molecular maps of diseases from omics data and network embeddings"

#### Dataset

| Disease | Omics | Positives | Negatives |
|---|---|---|---|
| Atopic dermatitis | GWAS and RNA-seq integration | 1000 | 15622 |
| Ulcerative colitis | GWAS and RNA-seq integration | 1300 | 14422 |
| Focal epilepsy | RNA-seq | 1106 | 18516 |
| Colorectal cancer | Somatic mutation | 82 | 19540 |
| Lymphoma | Somatic mutation | 86 | 19536 |
| Melanoma | Somatic mutation | 83 | 19539 |
| Aortic aneurysm | Proteomics | 109 | 1849 |

***Table S1. Training set size.*** *Dataset composition showing the number of positive and negative protein examples used to train logistic regression models for each of the seven diseases, along with the corresponding omics data type used.*

| Disease | Positives | Negatives |
|---|---|---|
| Atopic dermatitis | 188 | 188 |
| Ulcerative colitis | 98 | 98 |
| Focal epilepsy | 121 | 121 |
| Colorectal cancer | 49 | 49 |
| Lymphoma | 247 | 247 |
| Melanoma | 404 | 404 |
| Aortic aneurysm | 98 | 98 |

***Table S2. Test set size.*** *Composition of balanced literature-based gold standard test sets used for model evaluation, with equal numbers of positive and negative protein examples for each disease.*

| | | Jaccard Index |
|---|---|---|
| Atopic dermatitis | Ulcerative colitis | 0.06 |
| Colorectal cancer | Lymphoma | 0.07 |
| Colorectal cancer | Melanoma | 0.13 |
| Melanoma | Lymphoma | 0.09 |

***Table S3. Jaccard index of the training set.*** *Pairwise similarities between positive training protein sets for different diseases, showing low overlap (0.06-0.13) that validates the diseases have largely distinct associated protein sets despite potential shared mechanisms.*

### Cross-validation curves

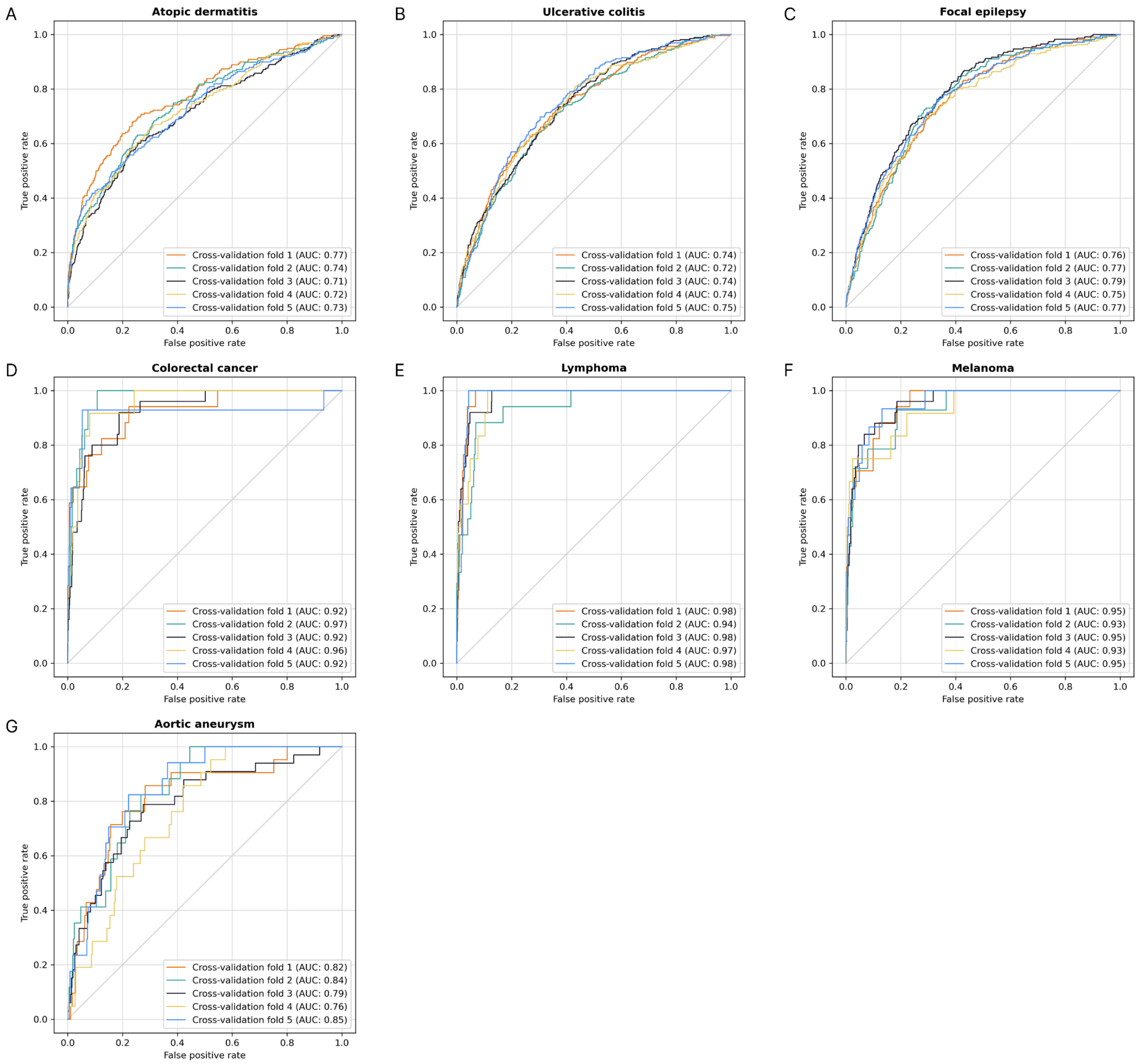


***Figure S1. Five-fold cross-validation of the network-based AI method.*** *(A) Atopic dermatitis. (B) Ulcerative colitis. (C) Focal epilepsy. (D) Colorectal cancer. (E) Lymphoma. (F) Melanoma. (G) Aortic aneurysm. ROC curves for each of the five cross-validation folds across all seven diseases (A-G), with AUC scores labeled for individual folds to demonstrate robustness of the network-based AI method.*

### DeLong test of the ROC curves

| Method 1 | Method 2 | P-value |
|---|---|---|
| Network-based AI (AUC: 0.95) | GWAS (AUC: 0.74) | $4.54\times10^{-13}$ |
| Network-based AI (AUC: 0.95) | RNA-seq (AUC: 0.64) | $7.39\times10^{-24}$ |
| Network-based AI (AUC: 0.95) | Integrated omics (AUC: 0.80) | $3.72\times10^{-09}$ |

***Table S4. DeLong test results on the atopic dermatitis literature-based gold standard.*** *Statistical comparison of ROC curve areas showing the network-based AI method significantly outperforms GWAS alone, RNA-seq alone, and integrated omics approaches ($P < 10^{-8}$).*

| Method 1 | Method 2 | P-value |
|---|---|---|
| Network-based AI (AUC: 0.95) | GWAS (AUC: 0.65) | $1{,}71\times10^{-25}$ |
| Network-based AI (AUC: 0.95) | RNA-seq (AUC: 0.69) | $1.50\times10^{-21}$ |
| Network-based AI (AUC: 0.95) | Integrated omics (AUC: 0.73) | $1.63\times10^{-17}$ |

***Table S5. DeLong test results on the ulcerative colitis literature-based gold standard.*** *Statistical comparison of ROC curve areas showing the network-based AI method significantly outperforms GWAS alone, RNA-seq alone, and integrated omics approaches ($P < 10^{-16}$).*

### Number of predicted disease-associated proteins

| Disease | Total proteins | Positive in omics data | Positive in text mining | Positive in both | Novel predcitions |
|---|---|---|---|---|---|
| Atopic dermatitis | 1717 | 370 | 157 | 68 | 1258 |
| Ulcerative colitis | 2647 | 448 | 160 | 50 | 2089 |
| Focal epilepsy | 1329 | 280 | 67 | 13 | 995 |
| Colorectal cancer | 2213 | 76 | 46 | 13 | 2104 |
| Lymphoma | 3183 | 85 | 266 | 53 | 2885 |
| Melanoma | 3320 | 81 | 236 | 12 | 3015 |
| Aortic aneurysm | 996 | 67 | 45 | 7 | 891 |

***Table S6. Number of predicted disease-associated proteins.*** *Breakdown of total predicted proteins by evidence source, showing proteins identified in omics data, text mining, both omics data and text mining, and novel predictions without prior evidence in each disease.*
